# Neural mechanisms underlying leg muscle responses during cervical transcutaneous spinal cord stimulation

**DOI:** 10.64898/2026.02.06.702847

**Authors:** Natalie Phelps, Rodolfo Keesey, Rachel Hawthorn, Carolyn Atkinson, Ismael Seáñez

## Abstract

Transcutaneous spinal cord stimulation (tSCS) of the cervical spinal cord has been thought to modulate lumbar networks, leading to the hypothesis that leg muscle recruitment may occur via recruitment of long-range spinal connections between cervical and lumbar circuits. To directly test this hypothesis, we compared arm and leg muscle responses elicited in unimpaired participants (N = 12) by cervical tSCS with the anodes placed over the iliac crests, with the anodes placed over the clavicles, and with lumbar tSCS as a control for leg muscle recruitment via the posterior root-muscle reflex. The idea of tSCS targeting cervico-lumbar connectivity would suggest that cervical stimulation could evoke responses in leg muscles. However, in our experiments, leg responses via cervical tSCS were only observed when the anodes were placed over the iliac crests, but not over the clavicles. These leg muscle responses had shorter latencies than those with lumbar tSCS and showed minimal post-activation depression, indicating efferent rather than afferent recruitment. Therefore, changes in leg muscle excitability by cervical-iliac tSCS previously attributed to descending cervical circuits could instead be explained by direct recruitment of efferent fibers near the iliac anodes. These findings suggest that cervical tSCS alone does not engage leg muscle motoneurons via long-range spinal or bidirectional pathways. Therefore, our study highlights the need to carefully consider electrode configuration when interpreting cervical tSCS mechanisms and additional or unexpected rehabilitative effects that extend caudally from the cervical spinal cord.

## Introduction

Transcutaneous spinal cord stimulation (tSCS) for the cervical region of the spinal cord has emerged as a promising neuromodulation technique to improve upper limb motor function of people with spinal cord injury (SCI)^1–4^. Although this technique is typically intended to target cervical spinal networks, multiple studies in cervical and multi-site tSCS have reported the modulation of lumbar networks^5–8^, leading to the hypothesis that leg muscle recruitment may occur via long-range spinal connections between cervical and lumbar networks^6,7,9–12^. However, the neural recruitment mechanisms for activation of lumbar networks via cervical tSCS remain unknown.

A defining feature of tSCS is its ability to activate large-diameter afferent fibers within the posterior roots^13–15^, enabling the interaction with supraspinal and spinal inputs^16–18^, which is thought to be essential for neuromodulation^18–21^. In contrast, activation of long-range propriospinal connections would be a previously unknown recruitment mechanism, suggesting that neuromodulatory effects of tSCS could extend to areas not directly targeted during tSCS-assisted rehabilitation, such as the arms in lumbar tSCS or the legs in cervical tSCS. Alternatively, direct activation of motor efferent fibers would suggest a recruitment mechanism similar to that in functional electrical stimulation, where action potentials are transmitted directly to the target muscle, bypassing spinal neural circuits^22^.

In this work, we investigate the neural basis of leg muscle activation during cervical tSCS by evaluating three potential mechanisms by which cervical stimulation could engage lumbar motor circuits (**Fig. 1a**). We propose that activation of leg muscles by cervical tSCS could be mediated by either i) activation of long-range propriospinal connections linking the cervical spinal cord and lumbar networks^9–12^ (**Fig. 1a,i**), ii) recruitment of efferent fibers along the lumbar plexus (**Fig. 1a,ii**), or iii) recruitment of afferent fibers in the lumbar spinal cord (**Fig. 1a,iii**).

**Figure 1.**
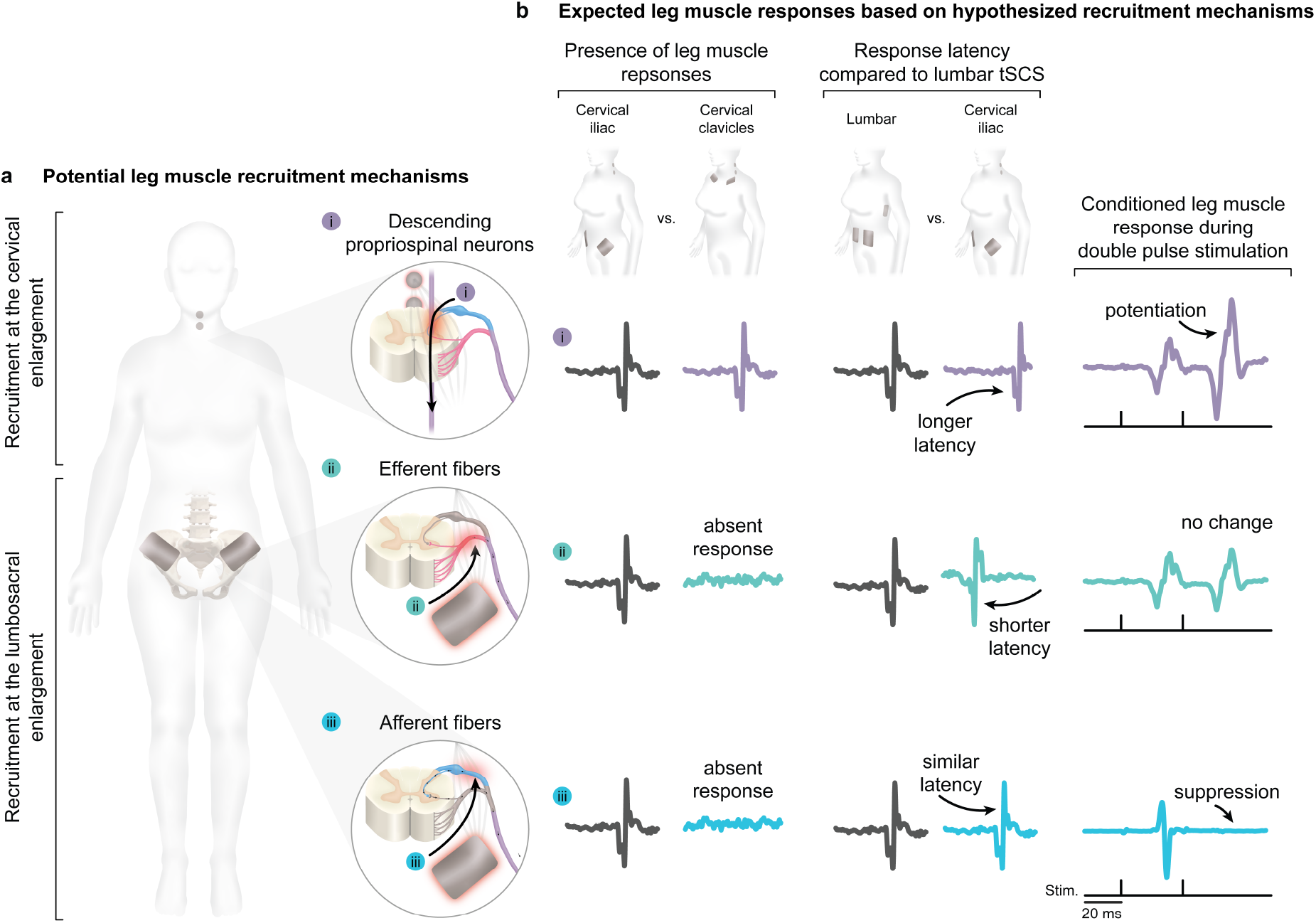
Hypothesized recruitment mechanism behind leg motoneuron activation by cervical transcutaneous spinal cord stimulation. **a**. Schematic of three recruitment mechanisms considered for activation of leg muscles via cervical tSCS. Leg responses by cervical tSCS could potentially be mediated by i) activation of long-range propriospinal connections linking the cervical spinal cord and lumbar networks, ii) recruitment of efferent fibers along the lumbar plexus, and iii) activation of afferent fibers in the lumbar spinal cord. **b**. Expected observations based on hypothesized recruitment mechanisms for activation of leg motoneurons. To elucidate the neural recruitment mechanisms for activation of leg muscles via cervical tSCS, we tested for the presence of leg muscle responses during cervical tSCS with the anodes over the iliac crests and the clavicles; we compared the leg muscle response latencies with cervical tSCS vs. those with lumbar tSCS; and we tested for modulation of the conditioned (second) response to double pulse stimulation, a marker of modulation at the synapse. Abbreviations: Stimulation (Stim.).

To test these hypothesized recruitment mechanisms, we compared arm and leg muscle responses elicited by cervical tSCS with two cathodes over the C3/C4 and C6/C7 vertebrae and the return anodes placed over the iliac crests (cervical-iliac)^3,4,23–26^, the anodes placed over the clavicles (cervical-clavicles), and with lumbar tSCS as a control for known leg muscle recruitment via the posterior root-muscle reflex (lumbar)^14,27–30^.

The cervical-iliac configuration was used as the primary condition for cervical tSCS, as it is commonly used to facilitate upper-extremity rehabilitation in people with SCI^3,4,23–25,31^. However, the placement of the return anodes over the iliac crests introduces the possibility that leg muscle recruitment results from activation of neural structures within the lumbosacral spinal cord or lumbar plexus, rather than from cervical cathodal stimulation. Although the cervical-clavicles configuration is also used in upper-extremity rehabilitation^4,24^, we included this as a control for cervical tSCS, as anodes are not present near the lumbosacral region in this configuration. Therefore, arm and leg muscle responses observed with the cervical-clavicles configuration would be more likely to originate from activation of neural circuits within the cervical region. Finally, we used lumbar-abdomen tSCS as a reference for the elicitation of posterior root-muscle reflexes by activation of afferent fibers in the lumbosacral spinal cord^14,27–30^.

Leg muscle responses via cervical tSCS were only observed when the anodes were placed over the iliac crests, but not over the clavicles. These leg muscle responses had shorter latencies than those with lumbar tSCS and showed minimal suppression due to post-activation depression, indicating efferent rather than afferent recruitment^14^. Therefore, we propose that activation of leg muscles by cervical tSCS can be explained by direct recruitment of efferent fibers near the return anodes over the iliac crests, rather than by activation of descending long-range propriospinal connections between the cervical and lumbar spinal cord.

## Methods

### Hypothesized neural recruitment mechanisms and their related observations

Three neural recruitment mechanisms were explored by comparing arm and/or leg muscle response properties under the three electrode configurations (**Fig. 1a**). We hypothesized that: i) If leg muscle responses are mediated by activation of cervical-to-lumbar long-range propriospinal or bidirectional connections, we would expect to observe leg muscle responses when the return electrodes are placed either over the iliac crests or over the clavicles. We would expect that leg muscle responses by cervical tSCS would have a longer latency than those with lumbar tSCS. Moreover, we would expect to observe a potentiation of the conditioned response during paired-pulse stimulation (**Fig. 1b,i)**. ii) If leg muscle responses are mediated by activation of efferent fibers at the lumbosacral enlargement, we would expect to observe a lack of leg muscle responses with cervical-clavicles tSCS. Moreover, we would expect leg muscle response latencies to be shorter than those with lumbar tSCS, and response amplitude to be unaffected by paired-pulse stimulation (**Fig. 1b,ii**). iii) If leg muscle responses are mediated by activation of the posterior roots, such as in lumbar tSCS, we would expect to observe a lack of leg muscle responses with cervical-clavicles tSCS, a leg muscle response latency close to that with lumbar tSCS, and a suppression of the conditioned response to paired-pulse stimulation, such as that in posterior root-muscle reflexes elicited by lumbar tSCS^14,27–30^ (**Fig. 1b,iii**).

### Participant recruitment

This study was reviewed and approved by the Washington University in St. Louis Institutional Review Board. Twelve unimpaired participants (7 female, 5 male, average age of 26.8 ± 3.99 years) were recruited and provided written informed consent prior to participation in this study. All twelve participants underwent all three electrode tSCS configuration conditions.

### Experimental conditions

To investigate recruitment mechanisms underlying leg activation during cervical tSCS, three electrode configurations were tested (**Fig. 2a**). In the cervical-iliac configuration, two circular surface electrodes (3.2 cm diameter, PALS Neurostimulation Electrodes, Axelgaard Manufacturing Co., Ltd., USA) were placed over the cervical spinal cord over the C3/C4 and C6/C7 vertebrae as cathodes, with two rectangular surface electrodes (7.5 × 10 cm) positioned bilaterally as anodes over the anterior iliac crests^3,4,23–26^. In the cervical-clavicles control configuration, two circular electrodes (3.2 cm diameter) were positioned at the C3/C4 and C6/C7 vertebrae as cathodes, and two rectangular electrodes (5 × 9 cm) were placed bilaterally over the clavicles as anodes^4,23,24,26^. The lumbar-abdomen configuration, known to preferentially recruit afferent pathways, involved placing a single rectangular electrode (5 × 9 cm) centered over the T11/T12 vertebrae as a cathode and two rectangular electrodes (7.5 × 10 cm) bilaterally over the abdomen as anodes^14,27–30^. Vertebral segments were identified by palpation following known anatomical landmarks^26,32^. Stimulation electrodes for all tSCS configurations were positioned prior to testing, and configurations were switched by re-connecting the electrode leads to match the appropriate electrode configuration. The order in which the electrode configurations were tested was randomized at the beginning of each experiment (randarray, Microsoft Excel).

**Figure 2.**
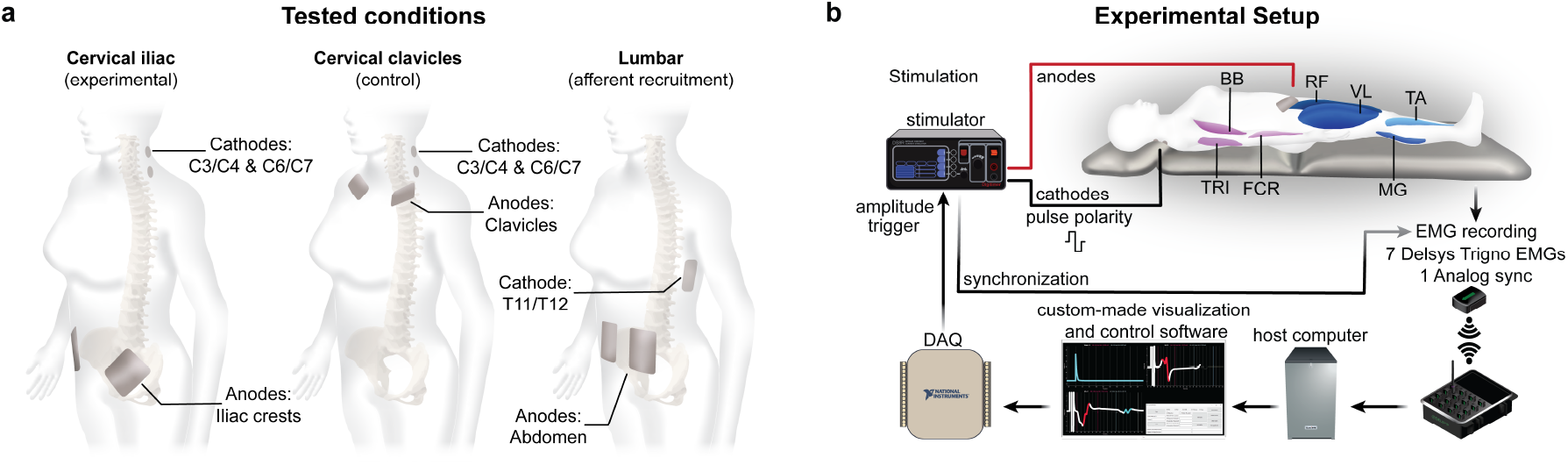
Experimental framework. **a**. The three tested cathode/anode electrode configurations for tSCS. **b**. Technological setup of tSCS experiments. Wireless EMG electrodes were used to record unilateral muscle activity from the upper and lower limbs. The data acquisition software that logged EMG signals additionally controlled stimulation onset and amplitude by the use of a microcontroller. More details in (*Data Acquisition*). Abbreviations: Rectus femoris (RF), vastus lateralis (VL), medial gastrocnemius (MG), tibialis anterior (TA), biceps brachii (BB), triceps (TRI), flexor carpi radialis (FCR).

### Experimental setup

Participants were positioned supine on a patient bed for the duration of the experiment (**Fig. 2b**). A pillow was placed under the participants’ heads for comfort, and their arms were relaxed at their sides. Surface electromyography (EMG) was recorded from seven muscles on the right side of the body using wireless sensors (Trigno® Avanti, Delsys Inc., USA). The muscles recorded were biceps brachii (BB), triceps long head (TRI), and flexor carpi radialis (FCR) in the upper limb, and the rectus femoris (RF), vastus lateralis (VL), medial gastrocnemius (MG), and tibialis anterior (TA) in the right lower limb. A Delsys analog to digital converter was used to synchronize EMG with the stimulation triggers for post-processing. EMG electrode placement followed SENIAM convention^33^.

### Data acquisition

In preparation for sensor placement, the skin was prepped with abrasive gel (NuPrep®, Weaver and Co. USA) and cleaned with alcohol pads. EMG signals were sampled at 4370 Hz using a custom-built software (Python 3.10.4) integrating the Trigno API and a National Instruments DAQ (USB-6001, National Instruments, USA) to visualize and record the data in real-time. Stimulation amplitudes were controlled by analog signals generated on the host computer and relayed to a National Instruments DAQ (USB-6001, National Instruments, USA), and pulse timing was relayed to a Raspberry Pi Pico microcontroller (Raspberry Pi Pico, Raspberry Pi Foundation, UK) before being transmitted to the National Instruments DAQ (**Fig. 2b**).

### Transcutaneous spinal cord stimulation

For each configuration, stimulation pulses were first manually delivered to identify the motor threshold and saturation/tolerance values. Motor threshold was determined by incrementally increasing stimulation amplitude until a muscle response amplitude greater than 50 µV was observed in either the arms or leg muscles for the cervical-iliac and cervical-clavicles configurations, or in the legs for the lumbar-abdomen^26^. Saturation was determined as the lowest stimulation amplitude at which all monitored muscles produced maximal peak-to-peak responses that no longer increased with further amplitude increments. If participants were not able to tolerate stimulation amplitudes to reach saturation, the maximum value was capped at the highest amplitude the participant could tolerate. These values were used to create recruitment curves for each electrode, which consisted of leg muscle response amplitudes across 16 stimulation amplitudes that were linearly spaced between the motor threshold and saturation/tolerance values, each repeated 3 times for a total of 48 pulses per condition. Stimulation was delivered in a descending amplitude order, which helped to enhance participant comfort and tolerance. Stimulation was delivered as 1 ms biphasic, paired pulses spaced 33 ms apart ^26,34^, with 7 seconds between paired pulses (with a randomized ±1 second interval to prevent stimulation anticipation), to allow for recovery cycles of posterior root-muscle reflexes^35^.

### Data analysis

EMG data processing was performed in MATLAB (Matlab 2020a, MathWorks, USA). EMG recordings were segmented relative to the first trigger pulse for each stimulation pulse. Three responses were averaged per stimulation amplitude, resulting in 16 averaged responses per muscle and electrode configuration. Peak-to-peak amplitudes were calculated as the difference between the maximum and minimum EMG values within the expected response time window. Responses were defined as present if they had peak-to-peak amplitudes of ≥ 50 µV.

#### Verification of evoked responses in leg muscles

We first aimed to determine whether leg muscle responses were present with the different electrode configurations. To test this, we visually inspected the average evoked responses across stimulation amplitudes and quantified whether the largest evoked response across participants was significantly greater than 50 µV (see *Statistics* for details).

#### Comparison of response latency to lumbar tSCS

To determine whether leg muscle responses by cervical tSCS were evoked rostrally or anteriorly to the posterior roots, we compared leg muscle response latencies with cervical-iliac tSCS to those with lumbar-abdomen tSCS. Response latency for responses at threshold was determined by visual inspection using a custom-made GUI in MATLAB and manually marking the response onset relative to the first stimulation pulse.

#### Verification of posterior root-muscle reflex

To determine whether leg muscle responses elicited via cervical-iliac tSCS were mediated by the posterior root-muscle reflex, such as those in lumbar tSCS, we quantified post-activation depression of the evoked responses with double-pulse stimulation at interstimulus intervals of 33 ms^26,34^.For each response, recruitment curves were generated as the muscle response peak-to-peak amplitudes as a function of stimulation amplitudes. Response suppression was quantified using the recruitment curves for the first and second muscle responses. The area under each curve (AUC_R1_, AUC_R2_) was computed using trapezoidal numerical integration (trapz, MATLAB). Suppression was then calculated for each muscle within each configuration using the following equation:

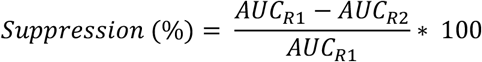

#### Motor Thresholds

We compared the ability of the different electrode configurations to elicit responses in leg muscles by comparing the motor thresholds with the cervical-iliac and lumbar-abdomen tSCS configurations. Additionally, we compared activation thresholds for arm muscles between the cervical-iliac and cervical-clavicles configurations.

## Statistical analysis

Group-level statistical analyses were implemented in Python (Python 3.11.4, Python Software Foundation, USA) using generalized linear mixed-effects models (GLMMs). Statistical comparisons (paired and one-sample *t*-tests) with a Bonferroni correction to account for multiple comparisons were performed in MATLAB (Matlab R2020a).

To evaluate whether leg and arm muscles responded to stimulation across all three electrode configurations, one-sample, one-tailed *t*-tests were conducted in MATLAB to test maximum muscle responses against 50 µV, with the null hypothesis that stimulation would not elicit responses greater than 50 µV (µ_₀_ ≤ 50 µV).

To determine whether recruitment of leg muscles by cervical-iliac tSCS occurred rostrally or anteriorly to the posterior roots, onset latencies were compared between cervical-iliac and lumbar-abdomen configurations. Two-tailed paired t-tests were performed, with the null hypothesis that response latencies between the two configurations would be equal (µ_d_ = 0).

To determine the degree of the response that was mediated by the recruitment of afferent fibers, such as in the posterior root-muscle reflex, we compared the amount of response suppression during paired-pulse stimulation between cervical-iliac and lumbar-abdomen tSCS. A GLMM with suppression as the dependent variable, configuration and muscles as fixed factors, and subjects as a random factor, was used to test for differences in suppression between electrode configurations and muscles. A one-sample *t*-test was used to test if suppression was significantly different from zero for arm and leg muscles. Paired-sample *t*-tests were used to test whether the amount of suppression was significantly different between responses elicited with the lumbar-abdomen vs. cervical-iliac configurations for each muscle under the null hypothesis that there would be no difference in suppression values between these configurations (µ_d_ = 0).

Additionally, we compared post-activation depression values between cervical-clavicles and cervical-iliac configurations to evaluate the degree of afferent recruitment in arm muscles. A one-sample two-tailed t-test was used to determine whether suppression values differed significantly from zero for each arm muscle. Paired-sample t-tests were then used to compare suppression magnitudes between cervical-clavicles and cervical-iliac tSCS for each muscle, with the null hypothesis that the two configurations would yield no difference in suppression (µ_d_ = 0).

To determine the minimal stimulation amplitudes needed to elicit leg muscle responses from each configuration, we compared motor thresholds between the cervical-iliac and lumbar-abdomen tSCS configurations. Two-tailed, paired-sample *t-*tests were performed to compare motor thresholds for leg muscles between the cervical-iliac and lumbar-abdomen tSCS conditions, under the null hypothesis that there would be no difference in motor thresholds between configurations (µ_d_ = 0).

## Results

### Leg muscle responses are elicited by cervical-iliac tSCS but not by cervical-clavicles tSCS

We first sought to confirm whether cervical tSCS elicits leg muscle responses and, if so, identify the source of recruitment (**Fig. 3a**). We considered two potential mechanisms of activation: i) activation originates from the cervical cathodes, with descending volleys traveling caudally through the spinal cord and synapsing on motoneurons at the lumbar enlargement to activate leg muscles, or ii) activation originates near the iliac crest anode electrodes. To test these alternative mechanisms, we first examined whether leg responses were present in both the cervical-iliac and cervical-clavicles configurations. If the activation of cervical-to-lumbar long-range propriospinal or bidirectional connections at the cervical level mediates leg muscle responses, we would expect to observe leg muscle responses with both the cervical-iliac and cervical-clavicles tSCS configurations. Alternatively, if the activation of spinal or peripheral networks along the lumbosacral enlargement mediates leg muscle responses, we would expect that only the cervical-iliac tSCS configuration would elicit these responses

**Figure 3.**
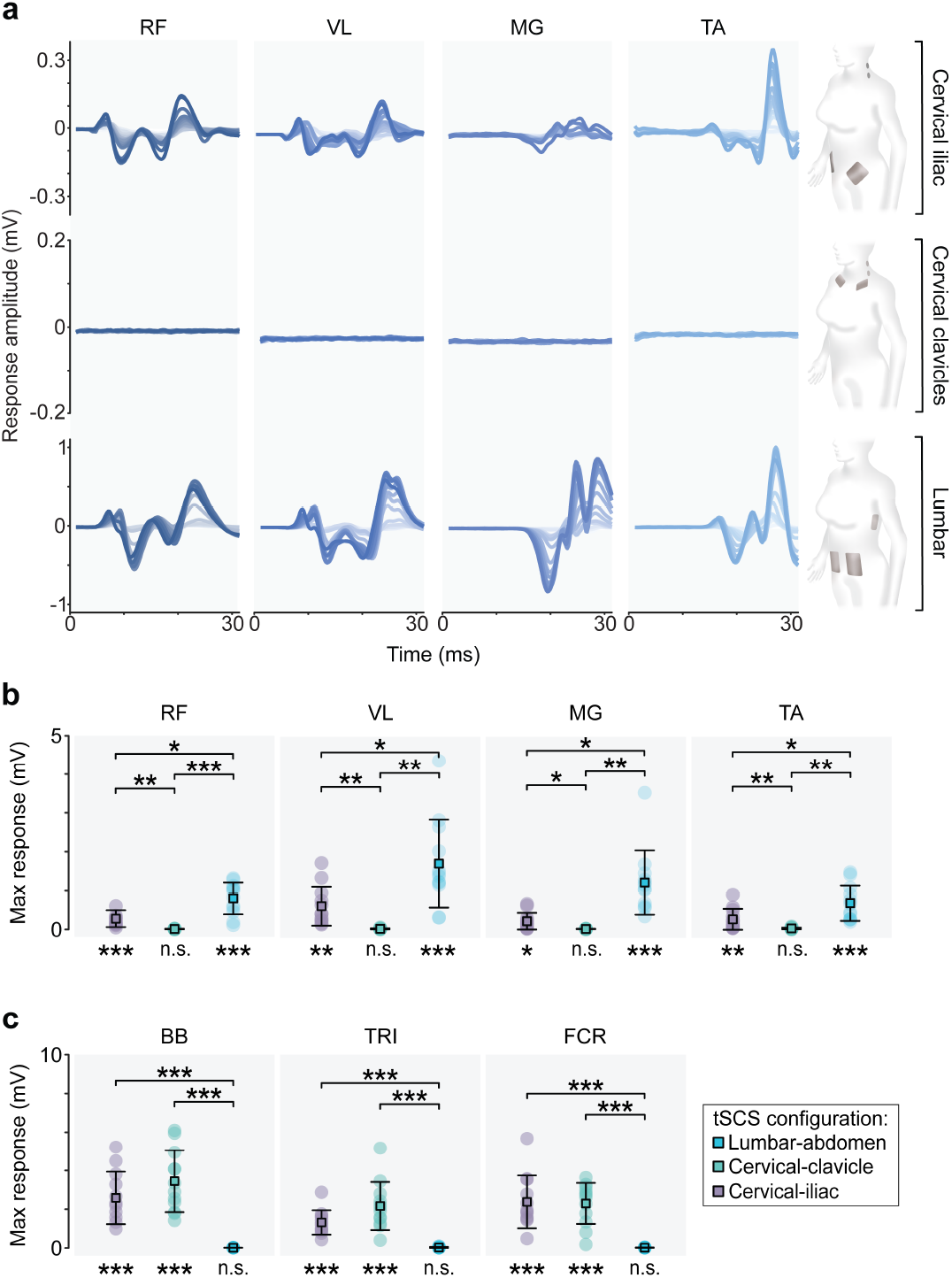
Leg muscles are activated during cervical tSCS with the anodes over the iliac crests, but not over the clavicles. **a**. Leg muscle responses from a representative participant (IA002). Three different electrode configurations were tested on the same day— cervical-iliac tSCS, cervical-clavicles tSCS, and lumbar-abdomen tSCS—across four leg muscles (RF, VL, MG, TA). Responses across the stimulation sweep are overlaid in gradient order with darker traces corresponding to higher stimulation amplitudes. Stimulation artifacts in the response waveforms were blanked for illustration purposes. **b**. Maximal responses in each leg muscle across configurations. **c**. Maximal responses in each arm muscle across configurations. Central boxes represent the estimated means, and whiskers indicate the 95% confidence interval. Asterisks below indicate the Bonferroni-corrected significance results for the one-sample *t*-tests of the maximum response of each muscle compared to 50 µV. Asterisks comparing two configurations above indicate the Bonferroni-corrected significance results for the paired *t*-tests of the maximum responses of each muscle compared across configurations. * p < 0.05; ** p < 0.01, *** p < 0.001. Abbreviations: Rectus femoris (RF), vastus lateralis (VL), medial gastrocnemius (MG), tibialis anterior (TA), biceps brachii (BB), triceps (TRI), flexor carpi radialis (FCR), millivolts (mV), milliseconds (ms), no significance (n.s.), transcutaneous spinal cord stimulation (tSCS).

To evaluate the presence of leg muscle activation during cervical tSCS, we tested whether the group-level maximal response of each muscle exceeded our predefined threshold of 50 µV via one-sample *t-*tests (**Fig. 3b**). In cervical-iliac tSCS, all four leg muscles exhibited group responses significantly greater than 50 µV (RF: t(11) = 3.651, p = 0.0076; VL: t(11) = 3.795, p = .0059; MG: t(11) = 2.627, p = 0.0471; TA: t(11) = 2.722, p = 0.0397). However, none of the leg muscles showed significant group responses during cervical-clavicles tSCS (RF: t(11) = -24.130, p = 1.000; VL: t(11) = -9.427, p = 1.000; MG: t(11) = -21.865, p = 1.000; TA: t(11) = -4.082, p = 1.000). In the lumbar-abdomen configuration, all four leg muscles exhibited group responses significantly greater than 50 µV (RF: t(11) = 6.335, p = 0.0001; VL: t(11) = 5.029, p = 0.0008; MG: t(11) = 4.850, p = 0.0010; TA: t(11) = 4.770, p = 0.0012).

To verify that adequate stimulation amplitudes for muscle recruitment were reached in cervical tSCS, arm muscle responses were also evaluated via one-sample *t-*tests (**Fig. 3c**). All three arm muscles exhibited responses significantly greater than 50 µV for cervical-clavicles tSCS (BB: t(11) = 7.357, p < 0.0001; TR: t(11) = 5.905, p = 0.0002; FCR: t(11) = 7.348, p < 0.0001) and cervical-iliac tSCS (BB: t(11) = 6.465, p < 0.0001; TR: t(11) = 6.994, p = 0.0002; FCR: t(11) = 5.911, p < 0.0001). Overall, none of the arm muscles showed significant group responses during lumbar-abdomen tSCS (BB: t(11) = -36.173, p = 1.000; TR: t(11) = -2.856, p = 1.000; FCR: t(11) = -10.844, p = 1.000).

Although leg muscle responses were not commonly elicited with the cervical-clavicles configuration, we did observe responses above 50 µV for one leg muscle (TA) in one participant (IA011), and these responses had latencies greater than 30 ms (**Supplementary Fig. 1a)**. Additionally, we observed responses above 50 µV in one arm muscle (TR) in one participant (IA006) from lumbar-abdomen tSCS with latencies over 40 ms (**Supplementary Fig. 1b**). In these two participants, additional muscles showed apparent responses by visual inspection. However, even at maximum tolerance, these were not above our pre-defined threshold of 50 µV.

The overall lack of leg responses in the cervical-clavicles condition (**Fig. 3a,b**) suggests that recruitment of cervical structures as a mechanism for elicitation of leg muscle responses with tSCS alone is unlikely. Instead, the presence of leg responses in cervical tSCS only when the return anodes are over the iliac crests suggests that recruitment occurs closer to neural structures within the lumbosacral enlargement.

### Leg muscle responses with cervical-iliac tSCS have shorter latencies than responses with lumbar-abdomen tSCS

To test whether leg muscle responses by cervical-iliac tSCS were elicited by activation of neural structures rostral to—or close to—the lumbar spinal cord, we compared their response latencies to those elicited by lumbar-abdomen tSCS (**Fig. 4a**). We considered three possible mechanisms of leg muscle activation during cervical-iliac tSCS: i) If recruitment occurs at the cervical spinal cord and propagates caudally through descending pathways to activate leg motoneurons, leg muscle response latencies with cervical-iliac tSCS would be substantially longer than those observed with lumbar-abdomen tSCS, due to the greater conduction distance; ii) if recruitment occurs along afferent fibers within the posterior roots, such as in lumbar-abdomen tSCS^26,28,34,36,37^, response latencies with cervical tSCS would be comparable to those with lumbar-abdomen tSCS; or iii) if recruitment occurs along efferent fibers within the anterior roots or lumbar plexus, response latencies with cervical-iliac tSCS would be shorter than those with lumbar-abdomen tSCS, as efferent activation would both start closer to the muscle (and have a shorter conduction distance) and bypass synaptic delay (typically 0.5 to 1 ms) inherent to afferent-driven reflexes^38,39^.

**Figure 4.**
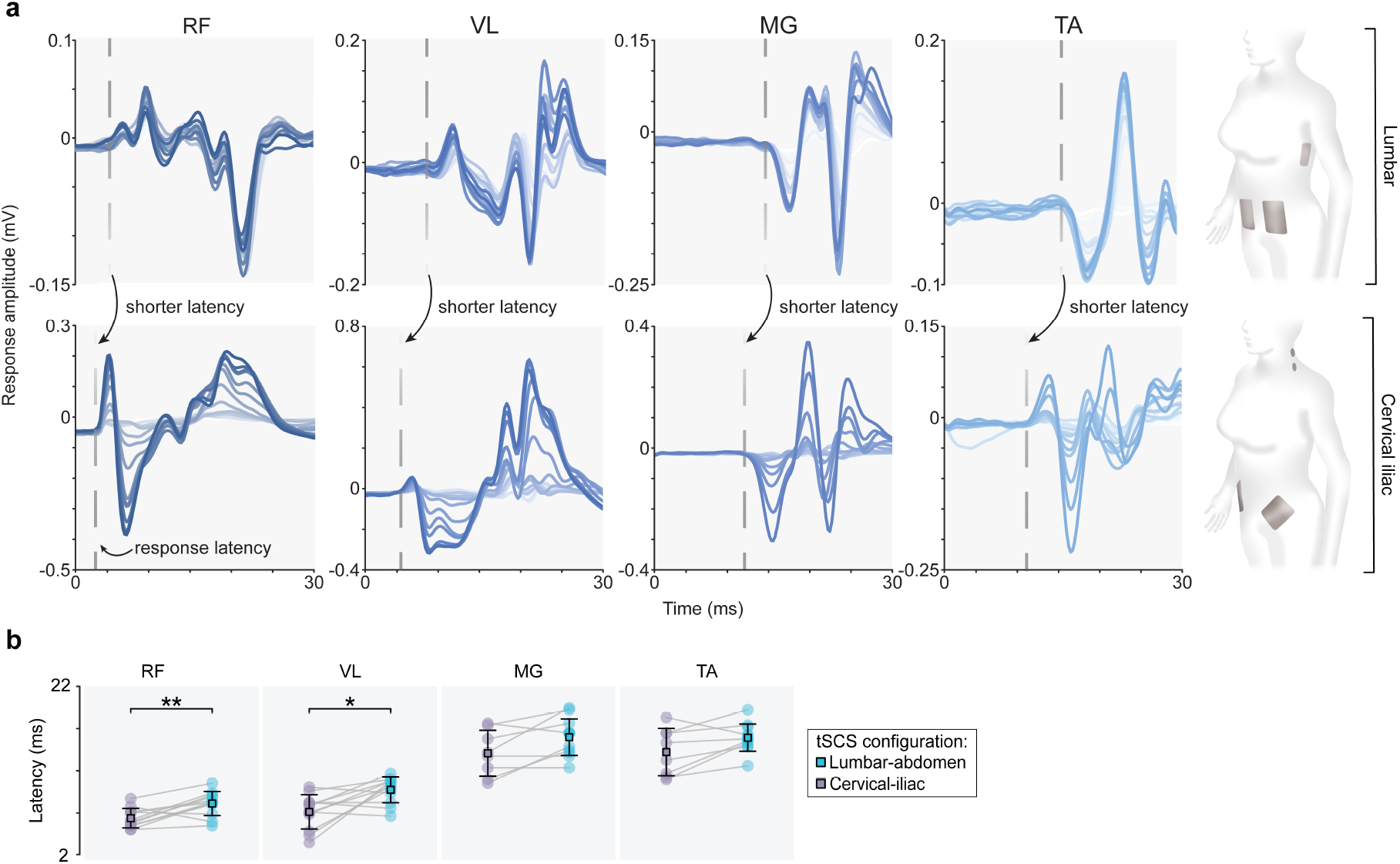
Comparisons of response latencies between cervical-iliac tSCS and lumbar-abdomen tSCS reveal differences in recruitment mechanisms. **a**. Leg muscle responses with lumbar and cervical-iliac tSCS in a representative participant (IA012). Dashed lines indicate response onset at the motor threshold, with time zero indicating stimulation onset. Note the leftward shift in response latency between lumbar and cervical tSCS. Responses across the stimulation sweep are overlaid in gradient order with darker traces corresponding to higher stimulation amplitudes. Stimulation artifacts were blanked for illustration purposes. **b**. Latency values for each leg muscle were compared between cervical-iliac tSCS and lumbar-abdomen tSCS at motor threshold. Latency was taken as the time between the stimulation onset and muscle response onset. Transparent circles represent individual participants. Central boxes represent the estimated means, and whiskers indicate the 95% confidence interval for each data group. Asterisks on top indicate the Bonferroni-corrected comparisons of paired t-tests between configurations within each muscle. * p < 0.05; ** p < 0.01. Abbreviations: Rectus femoris (RF), vastus lateralis (VL), medial gastrocnemius (MG), tibialis anterior (TA), millivolts (mV), milliseconds (ms), transcutaneous spinal cord stimulation (tSCS).

We observed that the proximal leg muscles (RF, VL) exhibited significantly shorter latencies during cervical-iliac tSCS compared to lumbar-abdomen tSCS (**Fig. 4b**), while distal muscles (MG, TA) showed no significant difference (RF: t(11) = -3.699, p = 0.0140; VL: t(11) = - 3.312, p = 0.0277; MG: t(7) = -2.422, p = 0.1840; TA: t(7) = -2.294, p = 0.2220). These findings suggest that cervical-iliac tSCS engages efferent pathways in proximal muscles, producing responses that originate closer to the muscle compared to lumbar tSCS. However, whether responses in distal muscles have a different origin than those in lumbar-tSCS remains unclear from this analysis alone. Nevertheless, as all leg responses evoked by cervical-iliac tSCS exhibited shorter or equal latencies than those elicited by lumbar-abdomen tSCS, activation of cervical pathways appears to be highly unlikely as the mechanism underlying the observed leg muscle responses.

### Leg muscle responses in cervical-iliac tSCS show poor evidence of post-activation depression in paired-pulse stimulation

To further investigate leg muscle recruitment mechanisms by cervical-iliac tSCS, we compared suppression due to post-activation depression during paired-pulse stimulation^26,34,36^ between the cervical-iliac and lumbar-abdomen tSCS configurations (**Fig. 5a,b**). The degree of afferent recruitment was evaluated by quantifying suppression of the second response relative to the first in the paired-pulse paradigm. The suppression of the second pulse is due to post-activation depression, which is mediated by synaptic mechanisms within the monosynaptic reflex pathway. This synaptic involvement signifies the activation of proprioceptive afferents, such as in lumbar tSCS^26,34,36^. In contrast, minimal or no suppression suggests direct activation of motor efferent fibers, which bypasses the synapse.

**Figure 5.**
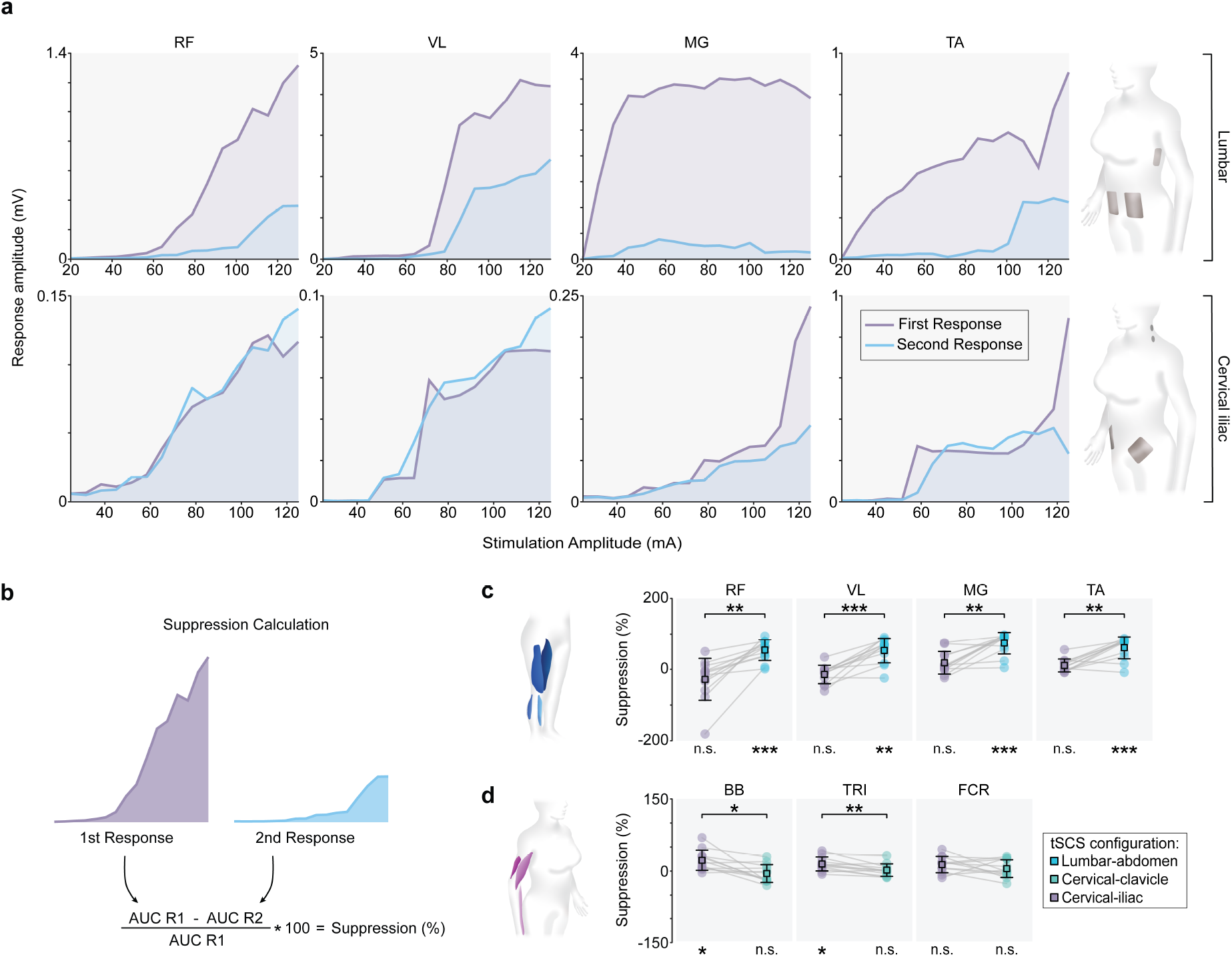
A high degree of post-activation depression in lumbar-abdomen tSCS but not in cervical-iliac tSCS suggests different recruitment mechanisms. **a**. Recruitment curves showing the peak-to-peak amplitudes of the first and second responses as a function of stimulation amplitude from one representative participant (IA003). Shaded regions indicate the area under each curve. **b**. Calculation of suppression using the area under the first and second recruitment curves. Suppression is calculated as a percent change from the first response. **c**. Suppression values for lumbar-abdomen and cervical-iliac tSCS in leg muscles across participants. Positive values reflect suppression, where the AUC of the second response was smaller than the AUC of the first response. **d**. Suppression values for cervical-clavicles and cervical-iliac tSCS in arm muscles across participants. Transparent circles represent individual participants, central boxes represent the estimated means, and whiskers indicate the 95% confidence interval for each data group. The asterisks below each group indicate the Bonferroni-corrected significance results for one-sample *t*-tests against zero. Asterisks on top indicate the Bonferroni-corrected comparisons of paired *t*-tests between configurations within each muscle. * p < 0.05, ** p < 0.01, *** p < 0.001. Abbreviations: Rectus femoris (RF), vastus lateralis (VL), medial gastrocnemius (MG), tibialis anterior (TA), biceps brachii (BB), triceps (TRI), flexor carpi radialis (FCR), no significance (n.s.), area under the curve (AUC), milliamperes (mA), millivolts (mV), transcutaneous spinal cord stimulation (tSCS).

Results from the Generalized Linear Mixed Model (GLMM) model (2 factors: Muscle [RF, VL, MG, TA]; Condition [cervical-iliac configuration vs. lumbar-abdomen configuration]) revealed a significant degree of suppression for the lumbar-abdomen configuration (β = 0.603, SE = 0.072, z = 8.380, p < 0.001), but not for the cervical-iliac configuration (β = -0.034, SE = 0.072, z = - 0.466, p = 0.641). One-sample *t*-tests (**Fig. 5c**) against zero showed that all four muscles under lumbar-abdomen tSCS exhibited a significant degree of suppression (RF: t(11) = 6.389, p = 0.0002; VL: t(11) = 5.258, p = 0.0011; MG: t(11) = 8.281, p < 0.0001; TA: t(11) = 6.706, p = 0.0001), whereas none of the muscles under cervical-iliac tSCS showed significant suppression (RF: t(11) = -1.659, p = 0.5017; VL: t(11) = -1.933, p = 0.3175; MG: t(11) = 2.005, p = 0.2808; TA: t(11) = 2.050, p = 0.2601). Further, paired *t-*tests between lumbar-abdomen and cervical-iliac configurations showed a significantly higher degree of suppression in all four leg muscles with lumbar-abdomen tSCS (RF: t(11) = -4.975, p = 0.0017; VL: t(11) = -5.808, p = 0.0004; MG: t(11) = -4.774, p = 0.0023; TA: t(11) = -5.090; p = 0.0014). These results suggest that while lumbar-abdomen tSCS primarily recruits afferent-mediated pathways^26,34,36^, cervical-iliac tSCS is more likely to recruit efferent fibers, bypassing synaptic transmission as in the posterior root-muscle reflex^14,18,40,41^.

To determine the degree of proprioceptive afferent recruitment in arm muscles between cervical-clavicles and cervical-iliac tSCS, we further compared suppression values in the paired-pulse paradigm (**Fig. 5d**). One sample *t*-tests against zero showed that two of the three arm muscles BB, TRI) under cervical-iliac tSCS displayed a significant degree of suppression (BB: t(11) = 3.728, p = 0.0100; TRI: t(11) = 3.535, p = 0.0140; FCR: t(11) = 2.762, p = 0.0555), whereas none of the muscles under cervical-clavicles tSCS showed significant suppression (BB: t(11) = - 0.936, p = 1.0000; TRI: t(11) = 0.529, p = 1.0000; FCR: t(11) = 0.978, p = 1.0000). Further, paired *t*-tests between cervical-clavicles and cervical-iliac configurations showed a significantly higher degree of suppression in two of the three arm muscles (BB, TRI) with cervical-iliac tSCS (BB: t(11) = 3.436, p = 0.01669, TRI: t(11) = 3.760, p = 0.0095; FCR: t(11) = 1.150, p = 0.8231). These findings suggest that recruitment of upper-limb muscles by cervical-iliac tSCS engages afferent-mediated pathways to a greater extent than cervical-clavicles tSCS. However, overall, the degree of suppression in arm muscles by cervical tSCS is markedly lower than that by lumbar tSCS in the lower limb (t(11) = -5.034, p < 0.001), in agreement with previously reported observations by our group^26^.

### No differences in motor thresholds between leg responses elicited via lumbar tSCS and cervical-iliac tSCS

We compared the motor thresholds of leg muscle responses between lumbar-abdomen and cervical-iliac tSCS (**Fig. 6b**). Paired *t*-tests showed that no muscles showed significant differences in motor thresholds between lumbar-abdomen and cervical-iliac tSCS (RF: t(11) = 0.742, p = 1.0000; VL: t(11) = -1.021, p = 1.0000; MG: t(7) = 1.783, p = 0.4712; TA: t(7) = -0.699, p = 1.0000). Additionally, we compared the motor thresholds of arm muscle responses between cervical-iliac and cervical-clavicles tSCS (**Fig. 6c**). Paired *t-*tests showed that no muscles showed significant differences in motor thresholds between cervical-iliac and lumbar-abdomen tSCS (BB: t(11) = 2.331, p = 0.1194; TRI: t(11) = 2.257, p = 0.136; FCR: t(11) = -1.105, p = 0.8777). The similarity in motor thresholds across configurations indicates that similar amounts of charge are necessary to elicit responses at threshold.

**Figure 6.**
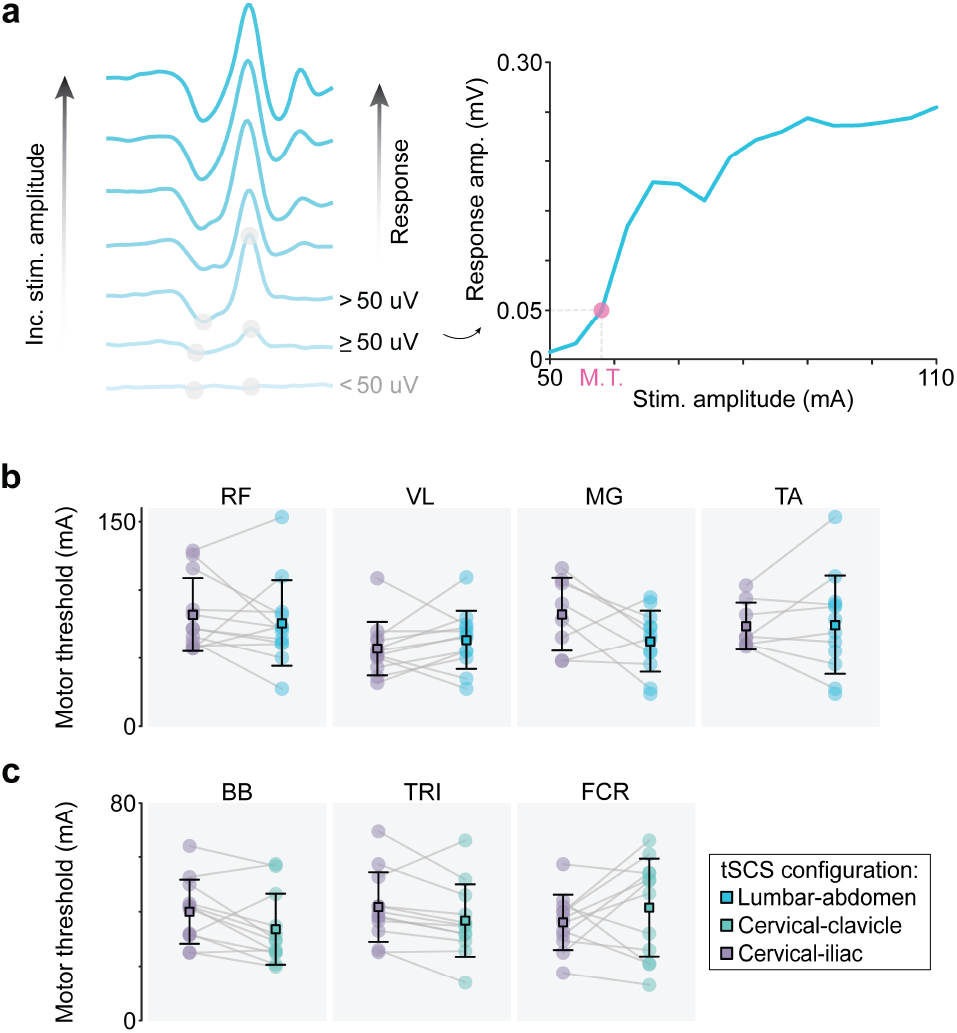
Leg muscles have similar motor thresholds with lumbar tSCS and cervical-iliac tSCS. **a**. Determination of motor threshold. Motor threshold is the lowest stimulation amplitude that evokes a muscle response with a peak-to-peak amplitude greater than or equal to 50 µV. **b**. Leg muscle motor thresholds compared across electrode configurations. **c**. Arm muscle motor thresholds compared across electrode configurations. Central boxes represent the estimated means, and whiskers indicate the 95% confidence interval for each data group. No significant differences were observed between configurations (Bonferroni-corrected paired *t*-test). Abbreviations: Rectus femoris (RF), vastus lateralis (VL), medial gastrocnemius (MG), tibialis anterior (TA), biceps brachii (BB), triceps (TRI), flexor carpi radialis (FCR), milliamperes (mA), millivolts (mV), stimulation (stim.), amplitude (amp.), increasing (inc.), transcutaneous spinal cord stimulation (tSCS), motor threshold (M.T.).

## Discussion

### Recruitment of leg motoneurons is likely to originate at the return electrode in cervical-iliac tSCS

Leg muscle responses were observed during both cervical-iliac and lumbar-abdomen tSCS, whereas cervical-clavicles tSCS failed to elicit detectable activation in the lower limbs. In our experimental design, the potential presence of leg responses during cervical-clavicles tSCS could have suggested recruitment through long-range propriospinal pathways linking cervical and lumbar segments, as all electrodes in this configuration were confined to the cervical region. Therefore, the absence of such responses suggests that stimulation of the cervical spinal cord alone is insufficient to engage lumbar motoneuron pools. This pattern implies that activation of leg muscles during cervical-iliac tSCS likely originated from local excitation near the iliac return electrodes rather than from descending activity initiated at the cervical cathodes. Because the iliac electrodes are positioned closer to the lumbar plexus, they may act as secondary stimulation sites, producing an effect that directly recruits motor efferent pathways in the lower limbs. Indeed, complementary work by our group and colleagues in high-resolution computational modelling demonstrated that cervical tSCS, with iliac crest anodes, directs current through the entire upper body, producing initiation sites of action potentials in both cervical and lumbar spinal roots and peripheral nerves^42^. The cervical-iliac tSCS configuration enables direct recruitment of lumbar axons via volume conduction, providing a mechanistic explanation for leg muscle responses during cervical–iliac tSCS. These findings support the conclusion that lower-limb activation arises from distal anode-driven peripheral or lumbar recruitment, rather than long-range propriospinal transmission from the cervical cord.

Interestingly, a single participant exhibited small but reproducible leg muscle responses during cervical-clavicles tSCS. These responses were of lower amplitude than those in cervical-iliac tSCS but occurred at longer latencies. This rare observation could reflect a single case of activation transmitted through descending volleys from the cervical spinal cord, which could explain the marked differences in amplitude and latency compared to those elicited with cervical-iliac tSCS. Therefore, while activation of long-range propriospinal pathways is not the dominant neural recruitment mechanism for leg motoneuron pool activation in cervical tSCS, it may be possible in rare or specific cases.

### Cervical-iliac tSCS engages local efferent rather than afferent pathways

Having established that the leg muscle responses observed during cervical tSCS most likely originate from local activation near the iliac anodes rather than from descending cervical pathways, it was important to determine which neural pathways were primarily being recruited. Differentiating between afferent- and efferent-mediated activation is critical for interpreting stimulation outcomes and refining effective experimental protocols, as afferent recruitment and its associated synaptic transmission play key roles in facilitating plasticity and functional recovery^18–21,28,43^. To identify the likely neural recruitment mechanisms, we analyzed two parameters—response latency and post-activation depression—each providing insight into whether stimulation engages afferent or efferent fibers ^26,34,36^.

We examined response latency as a proxy measure of the effective distance between the stimulation site and the recorded muscle. Afferent-mediated responses are expected to exhibit longer latencies than efferent-mediated ones due to their more proximal site of initiation within the reflex arc, as well as the time for the action potential to cross the synapse^38,39^. In contrast, efferent-mediated responses originate distal to these synapses, closer to the muscle, and therefore, reach the target muscle with shorter latencies. In our data, leg muscle responses during lumbar-abdomen tSCS exhibited significantly longer latencies than those evoked by cervical-iliac tSCS in proximal, but not distal muscles. Therefore, latency differences alone were not sufficient to distinguish between afferent and efferent recruitment for distal muscles.

One limitation of the latency analysis is that responses originating near the spinal cord may comprise a mixture of afferent- and efferent-mediated components^30^. Because onset latency is defined as the earliest time point at which the response exceeds a predetermined threshold, the measured latency primarily reflects the earliest activated component of the response. If an efferent-mediated component is present, it is likely to contribute to this initial rising phase and therefore dominate the latency estimate. Consequently, latency analysis can indicate the presence of an efferent-mediated contribution but cannot quantify the relative proportion of efferent vs. afferent recruitment within the overall response. For this reason, post-activation depression was employed as a complementary measure to more reliably distinguish between afferent- and efferent-mediated mechanisms of activation.

We employed a paired-pulse paradigm with a 33 ms interstimulus interval, as this duration has been shown to elicit measurable suppression in unimpaired individuals when afferent-mediated synaptic transmission is present^14,26,34^. Analysis of post-activation depression revealed distinct recruitment profiles across configurations. Lumbar-abdomen tSCS produced significant suppression in leg muscles across the recruitment range, whereas cervical-iliac tSCS showed no significant suppression. These results are in agreement with previous findings that lumbar-abdomen tSCS primarily activates proprioceptive afferent fibers, consistent with depletion of neurotransmitters due to synaptic transmission^44^. In contrast, the absence of suppression during cervical-iliac tSCS suggests direct recruitment of motor efferent fibers later in the reflex arc. Because these efferent fibers conduct action potentials directly to the muscle without the need for synaptic transmission, both pulses evoked via cervical-iliac tSCS elicit responses of comparable magnitude, resulting in little or no observable post-activation depression.

Motor threshold comparisons in the leg muscles revealed no significant differences between cervical-iliac and lumbar-abdomen tSCS. The comparable thresholds across configurations suggest that both activate neural structures with similar excitability, reflecting overlap in their recruited regions and engagement of neural networks with similar activation thresholds. Although these findings do not allow us to distinguish between afferent- and efferent-mediated mechanisms, the consistency in thresholds further supports the interpretation that cervical-iliac responses do not originate within the cervical spinal cord. If activation had occurred through descending cervical pathways, motor thresholds would be expected to differ substantially due to the markedly different neural structures being recruited^45–47^, together with the longer conduction distance and synaptic transmission involved. Instead, the similarity in thresholds implies recruitment of neural elements within a common lumbar region, consistent with local activation near the iliac return electrodes. This interpretation is consistent with biophysical models showing that posterior root afferents are preferentially engaged when current density peaks dorsally beneath a posterior cathode, whereas the large surface return electrodes produce more spatially diffuse current fields through the torso^48,49^. Positioned over the iliac crests, the return electrodes likely formed a broad anodal field where current exited the body, creating a secondary site of activation capable of directly depolarizing motor axons in the lumbar plexus—consistent with the efferent-dominant responses observed^48,49^.

### Potential for propriospinal facilitation during voluntary drive

Prior studies have shown that combining cervical tSCS with or without voluntary upper-limb movement, such as arm pedaling, enhances excitability in lower-limb muscles, suggesting that additional descending drive can facilitate propagation through these long spinal pathways^5–8^. However, these studies placed return electrodes over the iliac crests and included additional cathodes over the T11/L1 vertebrae. Therefore, changes in excitability of lower-limb muscles could potentially be attributed to these surface electrodes on the lower body. Nevertheless, one study using cervical-clavicles configuration showed that, with the right timing, cervical tSCS at suprathreshold intensity could facilitate the soleus H-reflex^13^.

Although leg muscle responses were not observed in the great majority of participants during cervical-clavicles tSCS, these findings do not definitively exclude the engagement of long-range propriospinal pathways linking cervical and lumbar regions during tSCS. Importantly, the present study was designed to isolate neurophysiological responses evoked in the absence of voluntary effort or overt movement. It therefore remains possible that cervical tSCS, when combined with voluntary activation, could recruit propriospinal circuits and modulate lumbar motoneuron pool excitability. Because relative motor thresholds varied substantially across participants and muscles, assessing responses at threshold during voluntary activation would have required complex, participant-specific experimental designs and additional control conditions. Such protocols were beyond the scope of the present study.

## Limitations

Prior studies have shown that lumbar tSCS with return electrodes over the abdomen consistently recruits afferent fibers, whereas configurations with returns over the iliac crests show limited afferent recruitment^29^. Therefore, although maintaining the anode over the iliac crests for lumbar tSCS would have removed the possibility of introducing a perceived confound, the lumbar-abdomen configuration was chosen as the reference for elicitation of posterior root-muscle reflexes^14,26,28–30,34^, rather than as a comparison between stimulation locations.

Surface stimulation introduces variability in current spread and depth of penetration, which can differ across individuals due to anatomical and tissue-conductivity differences^48,50^. While our results provide strong evidence for local efferent activation at the iliac return electrodes, computational modeling and high-resolution neuroimaging could further construct the precise current distributions underlying the observed responses^42,51^. Overall, these factors may be important to explore in future studies to refine our understanding of cervical tSCS mechanisms and their translational potential.

## Acknowledgements

All authors received partial support from the National Institutes of Health (NIH) NINDS Award Number K01NS127936, and internal funding from the Department of Biomedical Engineering, the Department of Neurosurgery, the McKelvey School of Engineering at Washington University in St. Louis. C.A. and R.H. received partial support from NIH NINDS Award Number T32NS126157.

The AI tool ChatGPT was used for language editing and grammar review during the preparation of this manuscript.

## Author contributions

N.P, R.K., and R.H. data analysis.

R.K., software development.

N.P. and R.K., conducted experiments.

I.S., conceptualization.

R.K. and I.S., supervision.

N.P., R.K., R.H., C.A., and I.S., data interpretation.

N.P. and I.S., manuscript writing, review, and editing.

All authors have reviewed and approved the manuscript.

## Supplementary Data

**Supplementary Figure 1.**
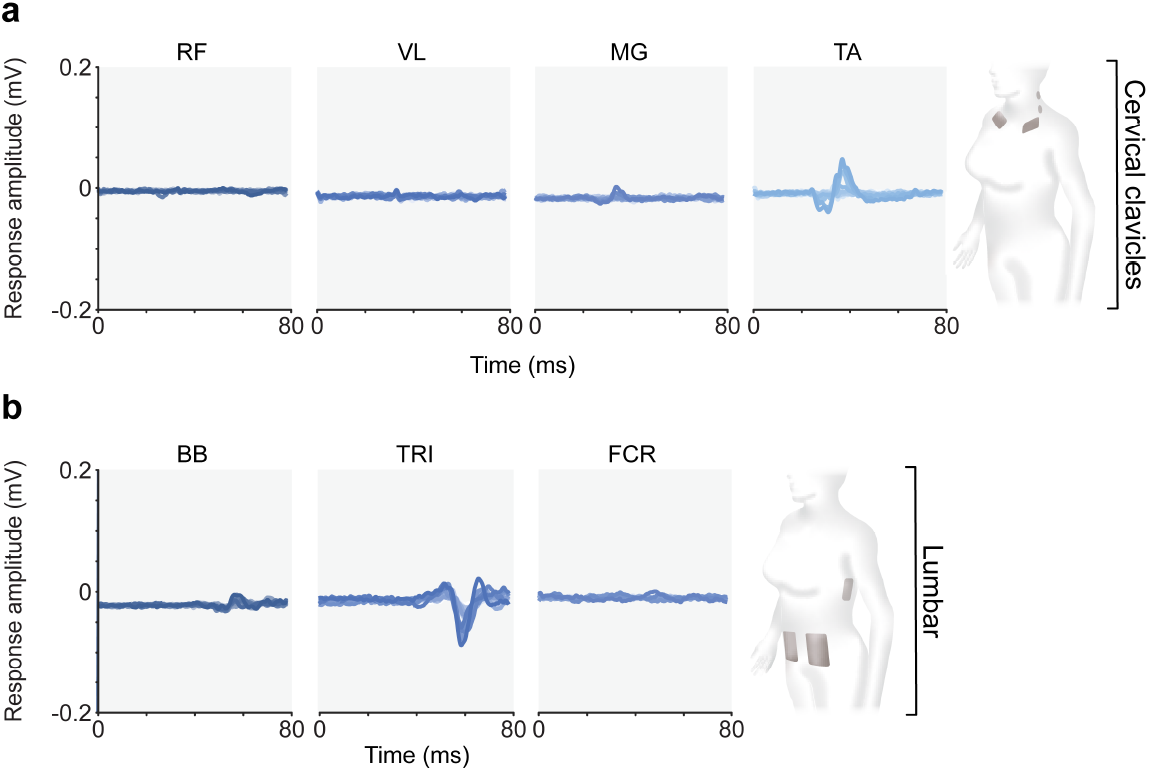
Single observations of responses in leg muscles during cervical tSCS and arm muscles during lumbar tSCS. **a**. Leg muscle responses from participant (IA011). Cervical-clavicle tSCS was administered up to the maximum stimulation amplitude tolerance, and responses were recorded across four leg muscles (RF, VL, MG, TA). This single participant exhibited a response (> 50 μV) in the TA from cervical stimulation. Although the other leg muscles appeared to have responses, these did not cross the threshold for detection. Time zero represents the time at which the first stimulation trigger was elicited. Responses overlaid with decreasing opacity on the same axes to illustrate recruitment across the 16 stimulation amplitudes. **b**. Arm muscle responses from participant (IA006). Lumbar-abdomen tSCS was administered, and responses were recorded across three arm muscles (BB, TRI, FCR). This single participant exhibited a response (> 50 μV) in the TRI from lumbar stimulation. Stimulation artifacts were blanked for illustration purposes. Abbreviations: Rectus femoris (RF), vastus lateralis (VL), medial gastrocnemius (MG), tibialis anterior (TA), biceps brachii (BB), triceps (TRI), flexor carpi radialis (FCR), millivolts (mV), milliseconds (ms), transcutaneous spinal cord stimulation (tSCS).

## Notes

### Competing Interest Statement

The authors have declared no competing interest.

## References

1. García-Alén, L. et al. Transcutaneous Cervical Spinal Cord Stimulation Combined with Robotic Exoskeleton Rehabilitation for the Upper Limbs in Subjects with Cervical SCI: Clinical Trial. Biomedicines 11, 589 (2023).

2. Chandrasekaran, S. et al. Targeted transcutaneous spinal cord stimulation promotes persistent recovery of upper limb strength and tactile sensation in spinal cord injury: a pilot study. Front. Neurosci. 17, (2023).

3. Inanici, F., Brighton, L. N., Samejima, S., Hofstetter, C. P. & Moritz, C. T. Transcutaneous Spinal Cord Stimulation Restores Hand and Arm Function After Spinal Cord Injury. IEEE Trans. Neural Syst. Rehabil. Eng. 29, 310–319 (2021).

4. Moritz, C. et al. Non-invasive spinal cord electrical stimulation for arm and hand function in chronic tetraplegia: a safety and efficacy trial. Nat. Med. 30, 1276–1283 (2024).

5. Fo, B., Ma, S. & S, U. Cervical Spinal Stimulation at Different Levels Evoked Multisegmental Motor Responses in the Lower Limbs. J. Case Rep. Stud. 7, 1 (2019).

6. Barss, T. S., Parhizi, B. & Mushahwar, V. K. Transcutaneous spinal cord stimulation of the cervical cord modulates lumbar networks. J. Neurophysiol. 123, 158–166 (2020).

7. Parhizi, B., Barss, T. S. & Mushahwar, V. K. Simultaneous Cervical and Lumbar Spinal Cord Stimulation Induces Facilitation of Both Spinal and Corticospinal Circuitry in Humans. Front. Neurosci. 15, 615103 (2021).

8. Samejima, S. et al. Multi-system benefits of non-invasive spinal cord stimulation following cervical spinal cord injury: a case study. Bioelectron. Med. 11, 20 (2025).

9. Huang, H. J. & Ferris, D. P. Upper and Lower Limb Muscle Activation Is Bidirectionally and Ipsilaterally Coupled. Med. Sci. Sports Exerc. 41, 1778–1789 (2009).

10. Zhou, R., Alvarado, L., Kim, S., Chong, S. L. & Mushahwar, V. K. Modulation of corticospinal input to the legs by arm and leg cycling in people with incomplete spinal cord injury. J. Neurophysiol. 118, 2507–2519 (2017).

11. Klarner, T. et al. Long-Term Plasticity in Reflex Excitability Induced by Five Weeks of Arm and Leg Cycling Training after Stroke. Brain Sci. 6, 54 (2016).

12. Ferris, D. P., Huang, H. J. & Kao, P.-C. Moving the Arms to Activate the Legs. Exerc. Sport Sci. Rev. 34, 113 (2006).

13. Islam, M. A. et al. Modulation of soleus H-reflex excitability following cervical transspinal conditioning stimulation in humans. Neurosci. Lett. 732, 135052 (2020).

14. Hofstoetter, U. S., Freundl, B., Binder, H. & Minassian, K. Recovery cycles of posterior root-muscle reflexes evoked by transcutaneous spinal cord stimulation and of the H reflex in individuals with intact and injured spinal cord. PloS One 14, e0227057 (2019).

15. Seáñez, I., Capogrosso, M., Minassian, K. & Wagner, F. B. Spinal Cord Stimulation to Enable Leg Motor Control and Walking in People with Spinal Cord Injury. in Neurorehabilitation Technology (eds Reinkensmeyer, D. J., Marchal-Crespo, L. & Dietz, V.) 369–400 (Springer International Publishing, Cham, 2022). doi:10.1007/978-3-031-08995-4_18.

16. Guiho, T., Baker, S. N. & Jackson, A. Epidural and transcutaneous spinal cord stimulation facilitates descending inputs to upper-limb motoneurons in monkeys. J. Neural Eng. 18, 046011 (2021).

17. Greiner, N. et al. Recruitment of upper-limb motoneurons with epidural electrical stimulation of the cervical spinal cord. Nat. Commun. 12, 435 (2021).

18. Minassian, K., McKay, W. B., Binder, H. & Hofstoetter, U. S. Targeting Lumbar Spinal Neural Circuitry by Epidural Stimulation to Restore Motor Function After Spinal Cord Injury. Neurother. J. Am. Soc. Exp. Neurother. 13, 284–294 (2016).

19. Holsheimer, J. Letters to the Editor. Neuromodulation Technol. Neural Interface 6, 270– 272 (2003).

20. Kathe, C. et al. The neurons that restore walking after paralysis. Nature 611, 540–547 (2022).

21. Formento, E. et al. Electrical spinal cord stimulation must preserve proprioception to enable locomotion in humans with spinal cord injury. Nat. Neurosci. 21, 1728–1741 (2018).

22. Kostyukov, A. I. et al. Fatigue effects in the cat gastrocnemius during frequency-modulated efferent stimulation. Neuroscience 97, 789–799 (2000).

23. Samejima, S. et al. Multisite Transcutaneous Spinal Stimulation for Walking and Autonomic Recovery in Motor-Incomplete Tetraplegia: A Single-Subject Design. Phys. Ther. 102, pzab228 (2022).

24. Freitas, R. M. de et al. Selectivity and excitability of upper-limb muscle activation during cervical transcutaneous spinal cord stimulation in humans. J. Appl. Physiol. https://doi.org/10.1152/japplphysiol.00132.2021 (2021) doi:10.1152/japplphysiol.00132.2021.

25. Keller, A. et al. Noninvasive spinal stimulation safely enables upright posture in children with spinal cord injury. Nat. Commun. 12, 5850 (2021).

26. Keesey, R. et al. Fundamental Limitations of Kilohertz-Frequency Carriers in Afferent Fiber Recruitment with Transcutaneous Spinal Cord Stimulation. 2024.07.26.603982 Preprint at 10.1101/2024.07.26.603982 (2024).

27. Hofstoetter, U. S. et al. Effects of transcutaneous spinal cord stimulation on voluntary locomotor activity in an incomplete spinal cord injured individual. Biomed. Tech. (Berl) 58 Suppl 1, /j/bmte.2013.58.issue-s1-A/bmt-2013-4014/bmt-2013-4014.xml (2013).

28. Hofstoetter, U. S., Freundl, B., Binder, H. & Minassian, K. Common neural structures activated by epidural and transcutaneous lumbar spinal cord stimulation: Elicitation of posterior root-muscle reflexes. PLOS ONE 13, e0192013 (2018).

29. Masugi, Y., Obata, H. & Nakazawa, K. Effects of anode position on the responses elicited by transcutaneous spinal cord stimulation. Annu. Int. Conf. IEEE Eng. Med. Biol. Soc. IEEE Eng. Med. Biol. Soc. Annu. Int. Conf. 2017, 1114–1117 (2017).

30. Minassian, K. et al. Posterior root-muscle reflexes elicited by transcutaneous stimulation of the human lumbosacral cord. Muscle Nerve 35, 327–336 (2007).

31. Gad, P. et al. Non-Invasive Activation of Cervical Spinal Networks after Severe Paralysis. J. Neurotrauma 35, 2145–2158 (2018).

32. Shin, S.Yoon, D.-M. & Yoon, K. B. Identification of the correct cervical level by palpation of spinous processes. Anesth. Analg. 112, 1232–1235 (2011).

33. Hermens, H. J., Freriks, B., Disselhorst-Klug, C. & Rau, G. Development of recommendations for SEMG sensors and sensor placement procedures. J. Electromyogr. Kinesiol. Off. J. Int. Soc. Electrophysiol. Kinesiol. 10, 361–374 (2000).

34. Bryson, N. et al. Enhanced selectivity of transcutaneous spinal cord stimulation by multielectrode configuration. BioRxiv Prepr. Serv. Biol. 2023.03.30.534835 (2023) doi:10.1101/2023.03.30.534835.

35. Hofstoetter, U. S., Freundl, B., Binder, H. & Minassian, K. Recovery cycles of posterior root-muscle reflexes evoked by transcutaneous spinal cord stimulation and of the H reflex in individuals with intact and injured spinal cord. PLOS ONE 14, e0227057 (2019).

36. Thatcher, K. L. et al. Optimizing transcutaneous spinal stimulation: excitability of evoked spinal reflexes is dependent on electrode montage. J. NeuroEngineering Rehabil. 22, 2 (2025).

37. Hofstoetter, U. S. et al. Augmentation of Voluntary Locomotor Activity by Transcutaneous Spinal Cord Stimulation in Motor-Incomplete Spinal Cord-Injured Individuals. Artif. Organs 39, E176–186 (2015).

38. Troni, W., Bianco, C., Coletti Moja, M. & Dotta, M. Improved methodology for lumbosacral nerve root stimulation. Muscle Nerve 19, 595–604 (1996).

39. Zhu, Y., Starr, A., Haldeman, S., Chu, J. K. & Sugerman, R. A. Soleus H-reflex to S1 nerve root stimulation. Electroencephalogr. Clin. Neurophysiol. 109, 10–14 (1998).

40. Oh, J. et al. Cervical transcutaneous spinal stimulation for spinal motor mapping. iScience 25, 105037 (2022).

41. de Freitas, R. M. et al. Selectivity and excitability of upper-limb muscle activation during cervical transcutaneous spinal cord stimulation in humans. J. Appl. Physiol. Bethesda Md 1985 131, 746–759 (2021).

42. Alashqar, A. et al. Virtual prototyping of non-invasive spinal cord electrical stimulation targeting upper limb motor function. 2026.01.22.701010 Preprint at 10.64898/2026.01.22.701010 (2026).

43. Barss, T. S., Parhizi, B., Porter, J. & Mushahwar, V. K. Neural Substrates of Transcutaneous Spinal Cord Stimulation: Neuromodulation across Multiple Segments of the Spinal Cord. J. Clin. Med. 11, 639 (2022).

44. Zucker, R. S. & Regehr, W. G. Short-Term Synaptic Plasticity. Annu. Rev. Physiol. 64, 355– 405 (2002).

45. Hofstoetter, U. S., Danner, S. M. & Minassian, K. Paraspinal Magnetic and Transcutaneous Electrical Stimulation. in Encyclopedia of Computational Neuroscience 1–21 (Springer, New York, NY, 2014). doi:10.1007/978-1-4614-7320-6_603-6.

46. de Freitas, R. M., Capogrosso, M., Nomura, T. & Milosevic, M. Optimizing sensory fiber activation during cervical transcutaneous spinal stimulation using different electrode configurations: A computational analysis. Artif. Organs 46, 2015–2026 (2022).

47. Minassian, K., McKay, W. B., Binder, H. & Hofstoetter, U. S. Targeting Lumbar Spinal Neural Circuitry by Epidural Stimulation to Restore Motor Function After Spinal Cord Injury. Neurotherapeutics 13, 284–294 (2016).

48. Danner, S. M., Hofstoetter, U. S., Ladenbauer, J., Rattay, F. & Minassian, K. Can the Human Lumbar Posterior Columns Be Stimulated by Transcutaneous Spinal Cord Stimulation? A Modeling Study. Artif. Organs 35, 257–262 (2011).

49. Ladenbauer, J., Minassian, K., Hofstoetter, U. S., Dimitrijevic, M. R. & Rattay, F. Stimulation of the human lumbar spinal cord with implanted and surface electrodes: a computer simulation study. IEEE Trans. Neural Syst. Rehabil. Eng. Publ. IEEE Eng. Med. Biol. Soc. 18, 637–645 (2010).

50. Holsheimer, J. Which Neuronal Elements are Activated Directly by Spinal Cord Stimulation. Neuromodulation J. Int. Neuromodulation Soc. 5, 25–31 (2002).

51. Guidetti, M. et al. Modeling Electric Fields in Transcutaneous Spinal Direct Current Stimulation: A Clinical Perspective. Biomedicines 11, 1283 (2023).

